# Unlocking hidden flaws in PPV and ACC: a step towards more reliable identification of protein complex

**DOI:** 10.1101/2025.03.03.641161

**Authors:** Yixiang Huang, Jiudong Wang, Xinqi Gong

## Abstract

As classic evaluation indexes, clustering-wise predictive positive value (*PPV*) and accuracy (*ACC*) have been widely used for the detection of protein complexes ([1]). However, we identified a critical error in their calculation, which can lead to inaccurate evaluation results under most conditions. Here, we elaborate on the problem of the original indexes and propose revised indexes *PPV*_*M*_ (*PPV* Modified) and *ACC*_*M*_, which correct the identified error. Experiments demonstrate that revised indexes achieve higher reliability. Based on the new indexes, we reevaluated three state-of-the-art computational methods for protein complex detection on five benchmarks to provide a revised baseline to facilitate comparison of performance for algorithms developed later. The code and data involved in the experimental section of this paper can be found in https://github.com/hyx-1/PPV_M-and-ACC_M.

## 1 Introduction

Since most intracellular biological processes are driven by protein complexes rather than individual proteins ([2]), recognition of protein complexes is of great importance. By understanding the composition and function of protein complexes, key molecular mechanisms, such as signal transduction, metabolic pathways, and regulation of gene expression, can be revealed ([3]). Computational methods play an indispensable role in protein complex detection, especially their efficiency and flexibility when processing large-scale data ([4]). By combining protein-protein interaction networks (PPI networks), structure prediction, and machine learning techniques, computational methods can rapidly screen candidate complexes to provide a priority list for experimental validation. This strategy not only saves experimental resources, but also reveals the composition of potentially unknown complexes and inspires drug design.

In the validation of computational methods, *PPV* and *ACC* are widely used as important indexes to evaluate the quality of the model ([5]). These two indexes were first proposed in the “Evaluation of Clustering Algorithms for Protein-protein Interaction Networks” in 2006 ([6]). Since being proposed, subsequent developed computational methods often compare these two indexes with state-of-the-art methods to demonstrate the superiority of their algorithms. By searching OpenAlex ([7]), the direct citations of this artical were as high as 862 by 2025.1.8. After further screening, *PPV* and *ACC* were used in 771 of these citations, accounting for 89%. We counted topics, the years of publication, types and citation counts of these citations (Fig. 1). It can be seen that these two indexes are widely used in bioinformatics, machine learning, and other fields (Fig. 1 (b) (c)), and have maintained high recognition and influence (Fig. 1 (d)).

**Fig 1.**
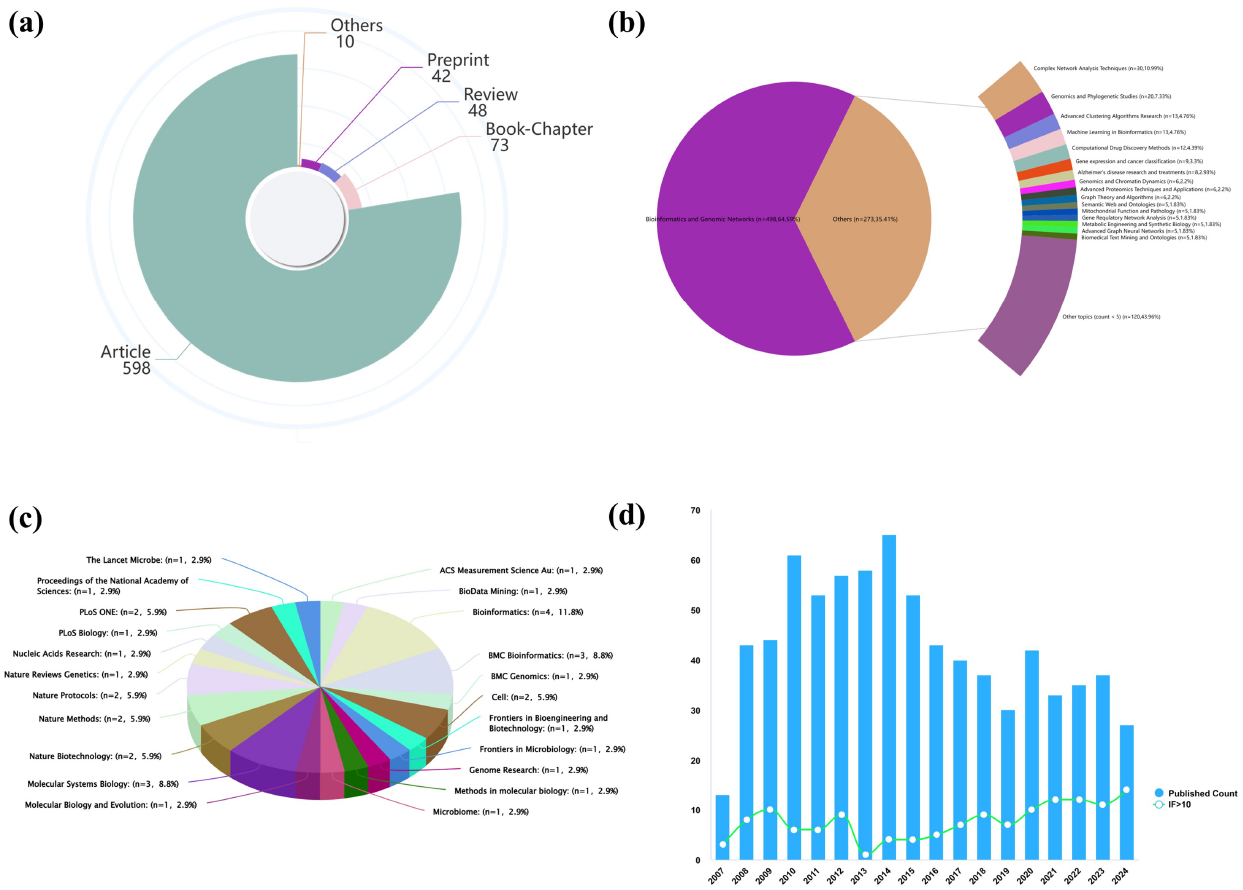
The information of the citations for “Evaluation of Clustering Algorithms for Protein-protein Interaction Networks”. (a): Types for citations that mention *PPV* and *ACC*. (b): Topics for citations that mention *PPV* and *ACC*. (c): Journal distributions for citations that mention *PPV* and *ACC* with more than two hundred citations. (d): Published Counts of all citations and citations with impact factor *>* 10 from 2007 to 2024 that mention *PPV* and *ACC*.

However, we found that *ACC* and *PPV* have serious problems and do not properly reflect the degree of similarity between the predicted protein complex set and the gold standard protein complex set. In the following, we elaborate theoretically on the problem of these indexes, supplemented by experimental proof. Finally, we propose new indexes *PPV*_*M*_ and *ACC*_*M*_ to evaluate computational methods for protein complex detection more accurately. We also verified the rationality of the new index through both theoretical and experimental aspects. In addition, we calculated *ACC*_*M*_ and *PPV*_*M*_ of three state-of-the-art methods on five benchmarks as baselines to facilitate the comparison of performance for algorithms developed later.

## 2 Results

### 2.1 Limitations and issues in *PPV* and *ACC*

*PPV* was defined as follows by Eq. 1.

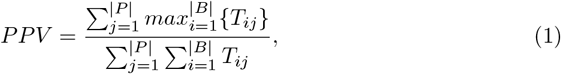

where *P* represents predicted set, *B* represents golden standard reference set, | · | denotes the protein complex number of the set and *T*_*ij*_ denotes the number of protein subunits discovered in both complex *i* from predicted set and complex *j* from reference set. ACC was calculated as follows:

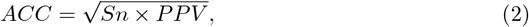

where the clustering-wise sensitivity (Sn) was determined by

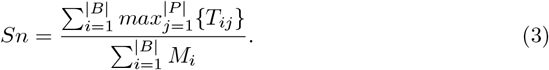

*M*_*i*_ denotes the number of proteins for complex *i* in the reference set. It is noted that the value of *ACC* partially depends on *PPV*, so we only discuss the problems and corrections of *PPV* later, and the correction of *ACC* is naturally the Eq. 4.

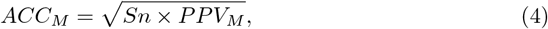

*PPV* aims to find the most similar protein complex in the reference set to each protein complex in the prediction set by maximum matching, and to describe the accuracy of the prediction set by considering similarity of each pair of matching protein complex. The idea of this index is feasible, but it ignores that certain proteins can be involved in the formation of multiple complexes. It treats protein complex detection as a community-separable clustering problem. In fact, the above problem is an overlapping community detection problem ([8]).

Let us examine the *l*-th protein complex of the prediction set, *PC*_*l*_, that is, fix *j* = *l* in Eq. 1. Observe that the numerator searches in the reference set for a complex with the largest intersection with *PC*_*l*_, which we denote as *RC*_*_. |*RC*_*_ ∩ *PC*_*l*_| will be added to the numerator term. At this time, if multiple complexes *RC*_1_, *RC*_2_, *RC*_*s*_ in reference sets have intersection with *PC*_*l*_ (*RC*_*_ ∈ {*RC*_1_, *RC*_2_ … *RC*_*s*_}), then the sum term in the denominator will increase | *RC*_1_ ∩ *PC*_*l*_ | + | *RC*_2_ ∩ *PC*_*l*_ | + … +| *RC*_*s*_ ∩ *PC*_*l*_|, which is usually much larger than the addition to the numerator. This situation makes it unreasonable that even if *RC*_*_ is in perfect agreement with *PC*_*l*_, a large *s* will make *PC*_*l*_ obtain a lower score.

In the following, we specify that the use of *PPV* as an evaluation index for protein complex detection has serious drawbacks. We believe that a qualified index should have the following properties:

1. The index should reflect the accuracy of the complex prediction set, and the index value should be consistent for prediction sets with the same accuracy. Here, *PPV* is the positive predictive value, which represents the fraction of proteins that truly belong to the same complex among all proteins predicted to belong to a complex. In this sense, assuming that the reference set is {126, 137, 145, 256}, then the *PPV* of the predicted sets {126, 137, 145, 256} and {137} should be the same and equal to 1 (Fig. 2(b)). However, after calculation by Eq. 1, the *PPV* is 0.5 and 0.6, respectively.

**Fig 2.**
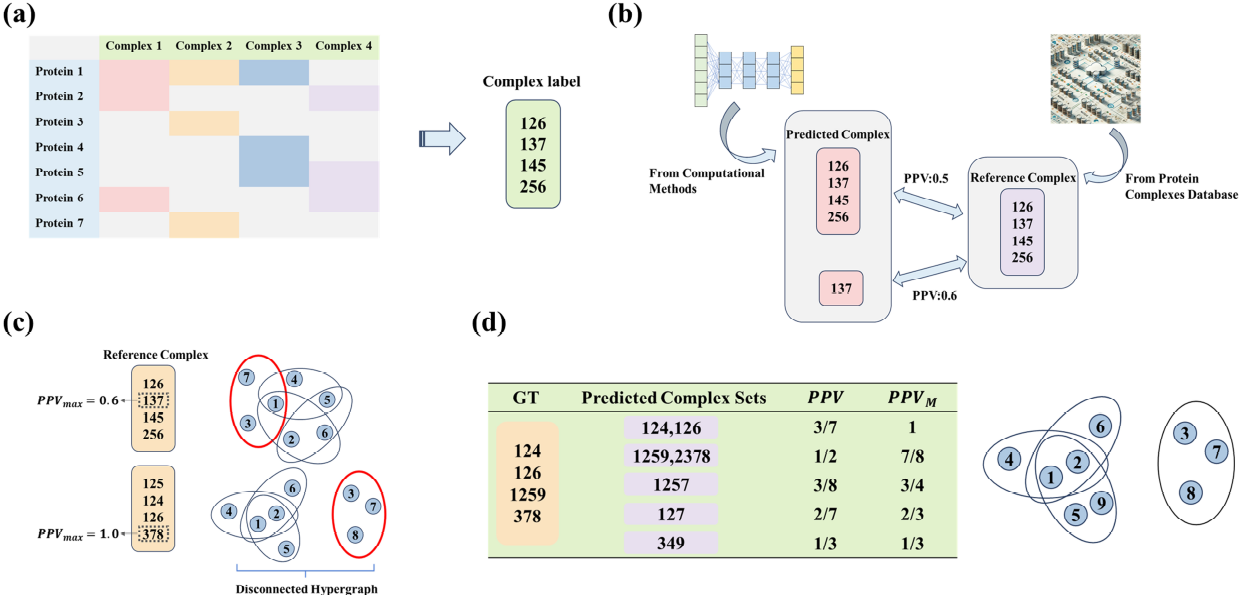
*PPV* and *PPV*_*M*_ in some cases. (a): Representation of protein complex composition. (b): *PPV* assigns a low score to the correct prediction set. (c): The *PPV* of different datasets has different upper bounds. (d): Comparison of *PPV* and *PPV*_*M*_ on the same prediction sets.

2. The index should have a fixed upper bound for different datasets and the index reaches its upper bound when the prediction set is the same as the reference set. Note that {126, 137, 145, 256} is the reference set (Fig. 2(b)), so the *PPV* of {126, 137, 145, 256} should be the largest, but 0.5 *<* 0.6 does not satisfy this. In addition, it can be seen from Fig. 2(c) that the maximum *PPV* corresponding to the two reference sets are 0.6 and 1.0 respectively, as proved in the Appendix. This means that when the first reference set is used as the gold standard complex set, no matter how accurate the predicted protein complex is, its *PPV* cannot reach a value above 0.6. It is easy to see that the reason for this phenomenon is that all the complexes in the reference set have at least one protein involved in forming the other complexes, so this part of the proteins repeatedly participates in the summation in the denominator. Only when the hypergraph with proteins as nodes and complexes as hyperedges is not connected, the *PPV* can reach a maximum of 1.0.

### 2.2 Corrected indexes for identification of protein functional modules: *PPV*_*M*_ and *ACC*_*M*_

In view of the above problems, we propose the new index *PPV*_*M*_ as a correction of *PPV*, which is calculated as follows:

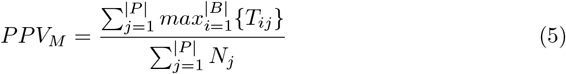

where *N*_*j*_ denotes the number of subunits contained in the *j*-th complex in the prediction set. It is not difficult to verify that *PPV*_*M*_ takes the value 1 in both cases of Fig. 2(b), the upper bound is 1 in both reference sets of Fig. 2(c), and the value is 1 when the predicted and reference sets are the same. Thus *PPV*_*M*_ satisfies the above two properties.

We further illustrate the superiority of *PPV*_*M*_ with some examples. For some given prediction sets, we compare their *PPV* with *PPV*_*M*_. The reference set is {124, 126, 1259, 378}, as shown in Fig. 2(d). When the prediction set is {124, 126}, both protein complexes are predicted correctly. In this case, *PPV*_*M*_ is 1, which reliably describes the accuracy of the prediction set, while *PPV* is only 3/8. When the prediction sets are {1259, 2378} and {1257}, the predicted protein complex is only one protein subunit away from the gold standard, and the positive predictive value should be close to 1 at this time. Note that *PPV*_*M*_ takes the values of 7/8 and 3/4, which can effectively characterize the positive predictive value, while *PPV* is only 1/2 and 3/8. In addition, for the {127} and {349} prediction sets, the former finds the core of 12, while the subunits of latter from three different complexes in the gold standard, so the positive predictive value of the former should be higher than that of the latter. The *PPV*_*M*_ value of the former is 2/3, and the latter is 1/3, which accurately distinguishes the different accuracy between them. However, the *PPV* values are 2/7 and 1/3, respectively, and the latter is instead larger, which is contrary to the real situation.

## 3 Experiments

### 3.1 Methods and materials

We selected ClusterOne, PC2P and AdaPPI, three classical protein complex detection methods, to verify the effectiveness of *PPV*_*M*_ and *ACC*_*M*_. ClusterONE ([9]) optimize the node scoring function by introducing the concept of overlapping clusters, thereby effectively coping with the challenge of proteins existing in multiple functional modules at the same time. PC2P ([10]) divides the network into subgraphs spanned by two shapes by removing the minimum number of edges in a given PPI network, and then solves the coherent partitioning problem using a parameter-free greedy approximation algorithm. AdaPPI ([11]) proposes a graph representation method that integrates protein interaction networks and GO annotation information, and then predicts protein functional modules through an adaptive graph convolutional network.

Considering the availability and size of the datasets, we selected five yeast PPI datasets containing different numbers of proteins to evaluate the index performance, including Collins ([12]), Krogan-core ([13]), Krogan14k ([13]), Biogrid ([14]), and the Database of Interacting Proteins (DIP [15]). These datasets are extremely common in protein complex detection algorithms, so it makes sense to show their *PPV*_*M*_ and *ACC*_*M*_. The details of the datasets are given in the Tab. 1. Additionally, we utilized the known protein complexes containing three or more proteins included in the HyperGraphComplex ([16]) as golden standard reference set, specifically including the MIPS ([17]), CYC2008 ([18]), Saccharomyces Genome Database (SGD) ([19]), Aloy ([20]) and TAP06 ([21]).

**Table 1.** Datasets information

| Datasets | Protein nodes | Interaction edges |
| --- | --- | --- |
| Collins | 1622 | 9074 |
| Krogan-core | 2708 | 7123 |
| Krogan14k | 3581 | 14076 |
| DIP | 4928 | 17201 |
| Biogrid | 5640 | 59748 |

It is easy to see from Eq. 1-5 that *PPV* (*PPV*_*M*_) measures how close the complex in the predicted set is to the reference complex, and *Sn* measures how much the complex in the reference set is predicted. *ACC*(*ACC*_*M*_) combines the two. Therefore, we use the index *Precision*, which also focuses on the accuracy of the prediction set itself, and the comprehensive index *F* 1 as references ([22]). Normally, *PPV* should have a strong positive correlation with *Precision*, while *ACC* has a strong positive correlation with *F* 1, which combines *Precision* with the index *Recall* that focuses on the coverage of the reference set ([1]). *Precision, Recall*, and *F* 1 are defined as follows:

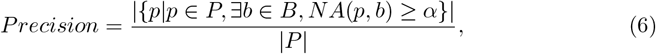

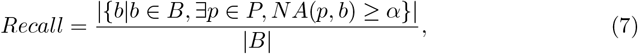

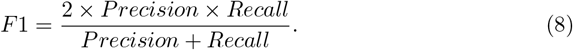

Where *α* is set to 0.25 according to [23], *NA* is defined as follows:

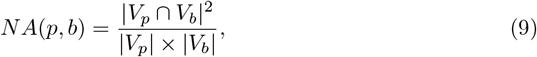

where *V*_*p*_ and *V*_*b*_ represent the sets composed by subunits of the protein complexes *p* and *b*.

### 3.2 Performance and advantages of *PPV*_*M*_ and *ACC*_*M*_

For index *X* and index *Y*, we use Pearson’s correlation coefficient to characterize their correlation degree in the *k*-th dataset:

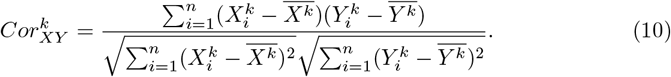

Among them, 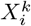 and 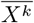 denote the value of the *i* algorithm in the *X* index on the *k*-th dataset and the average of all algorithms on the *k*-th dataset with respect to the *X* index. Same thing for 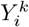 and 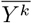.

We divide the correlation cases into three types according to the correlation coefficient: positive correlation, negative correlation and non-significant correlation as Eq. 11.

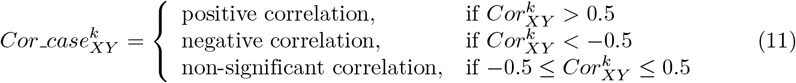

For the indexes with and without improvement, the correlation cases among the six indexes are shown in Fig. 3. Each pie chart contains a summary of the correlation case between two indexes across all datasets. The diagonal shows the correlation of each index with itself, which is clearly positive.

**Fig 3.**
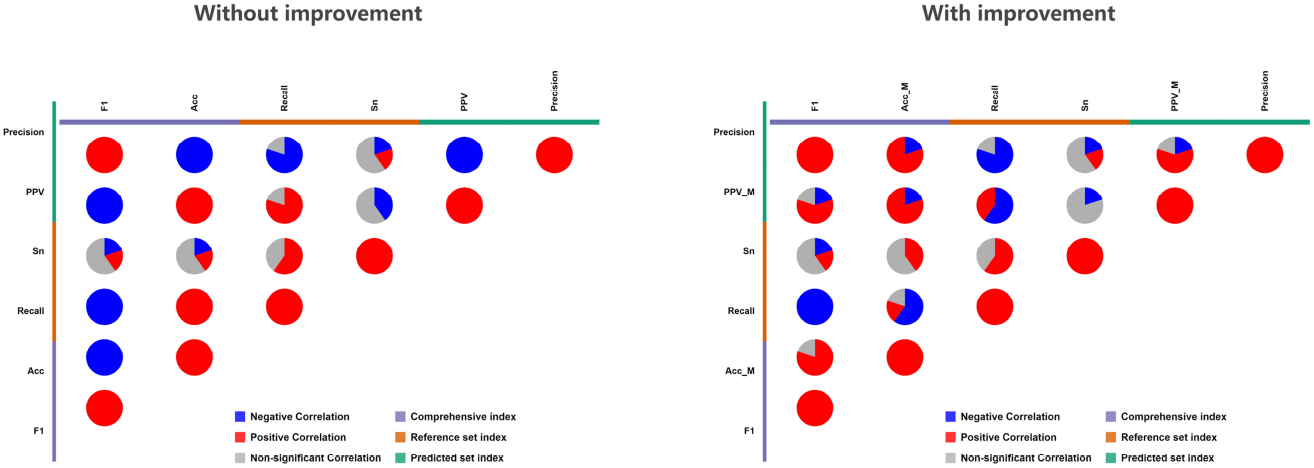
Correlation between indexes w/ and w/o improvement.

Observing Fig. 3, it can be seen that the reference set indexes *Recall* and *Sn* show a positive correlation overall, which is consistent with the reality. What changed before and after improvement were the predicted set index and the comprehensive index. *Precision* and *PPV* show a negative correlation on all datasets, while *Precision* and *PPV*_*M*_ show a positive correlation on most datasets. *F* 1 is negatively correlated with *ACC* on all datasets, while it is positively correlated with *ACC*_*M*_ on 80% of datasets. This indicates that the improvement of our proposed indexes *ACC*_*M*_ and *PPV*_*M*_ is reasonable, and the improved indexes have stronger evaluation ability for protein complex detection algorithms.

In addition, we display *ACC*_*M*_ and *PPV*_*M*_ of these three state-of-the-art methods on five benchmarks as baselines to facilitate the comparison of performance for algorithms developed later. Please see Tab. 2 and 3.

**Table 2.** Performance of different methods in *PPV*_*M*_ on the considered datasets.

| Methods | Datasets |  |  |  |  |
| --- | --- | --- | --- | --- | --- |
|  | Biogrid | Collins | DIP | Krogan14k | Krogan-core |
| Clusterone | 0.4859 | 0.5539 | 0.4870 | 0.8334 | 0.5008 |
| PC2P | - | 0.4624 | 0.2547 | 0.2463 | 0.2914 |
| AdaPPI | 0.3727 | 0.3806 | 0.4729 | 0.4120 | 0.5354 |

**Table 3.** Performance of different methods in *ACC*_*M*_ on the considered datasets.

| Methods | Datasets |  |  |  |  |
| --- | --- | --- | --- | --- | --- |
|  | Biogrid | Collins | DIP | Krogan14k | Krogan-core |
| Clusterone | 0.4731 | 0.6590 | 0.4346 | 0.4897 | 0.5391 |
| PC2P | - | 0.6370 | 0.3529 | 0.3793 | 0.4523 |
| AdaPPI | 0.5066 | 0.5474 | 0.5321 | 0.5301 | 0.6303 |

### 3.3 Illustration of the algorithm reproduction

We reproduced all three algorithms with default parameters, and the output was a list of protein complexes. In particular, for ClusterONE, we run the program using java software on Windows system. For AdaPPI, a deep learning algorithm that outputs a list of protein complexes in each training round, we use the output after all 500 rounds as the final result. For PC2P, we run in sequential mode. Due to its high time complexity, the Biogrid dataset took more than two months to run, so this part of the results is vacant. We run the two algorithms AdaPPI and PC2P using Ubuntu 18.04.6 LTS with an Intel(R) Xeon(R) CPU E5-2678 v3 @ 2.50GHz. See https://github.com/hyx-1/PPV_M-and-ACC_M for the detailed reproduction process.

## 4 Discussion

At this point in this article, careful readers have probably noticed that just as *Precision* and *Recall* have symmetries about prediction sets and reference sets, *Sn* and *PPV*_*M*_ also have certain symmetries, which is in fact where our inspiration comes from. Another impetus for this article is that we are developing a protein complex detection algorithm and found that *PPV* and *ACC* are always abnormally low when testing algorithm performance (laughter).

In addition, we observe an interesting phenomenon in Fig. 3. *Precision* and *Recall* show a negative correlation on most datasets, indicating that an algorithm is often unable to achieve both “accurate” and “complete”, that is, unable to take into account the accuracy of the prediction complex itself and the coverage of the reference set. Meanwhile, *F* 1 scores were positively correlated with *Precision* and negatively correlated with *Recall* in all five datasets in Fig. 3. It is observed that the positions of the two are exactly the same in Eq. 8, which means that moderately reducing the *Recall* of the model within an adjustable range can greatly improve the *Precision*. Therefore, how to fundamentally improve the *Recall* of protein complex detection becomes an important issue.

## 5 Conclusion

*PPV* and *ACC* are widely used indexes, and their accuracy and applicability directly affect the evaluation results of model performance. However, the existing *PPV* and *ACC* indexes can not properly reflect the degree of similarity between the prediction complex set and the gold standard complex set, which may lead to the deviation of the evaluation results, and thus affect the direction of model selection and optimization. We proposed and validated improved indexes *PPV*_*M*_ and *ACC*_*M*_, which can evaluate the model performance more comprehensively and objectively, and lay a foundation for developing better protein complex prediction methods. The improved *PPV*_*M*_ and *ACC*_*M*_ can not only be used for protein complex prediction, but may also be applied in other scenarios where the prediction results need to be evaluated for similarity to true values, such as gene regulatory network prediction or metabolic pathway identification. A theoretically solid and proven new index has the potential to be applied across fields.

## 6 Data and code availability

The code and data involved in the experimental section of this paper can be found in https://github.com/hyx-1/PPV_M-and-ACC_M. If you have any questions, please contact the following.

## 7 Appendix

### Claim 1.

*The maximum values of PPV for two reference sets in Fig. 2(c) are 0*.*6 and 1*.*0*.

*Proof*. For the second reference set, it is easy to verify *PPV*_{378}_ = 1. It follows from Eq. 1 that *PPV* ≤ 1 and hence *PPV*_*max*_ = *PPV*_{378}_ = 1. Next we just need to prove that the first reference set satisfies the claim. The protein complex detection problem considers only complexes containing at least three subunits. First we consider the case when the prediction set is a single complex. This can be divided into the following two cases:

1. The complex contains three subunits. Obviously, PPV has a chance to maximize when the prediction set is one of the reference set complexes. By calculation, 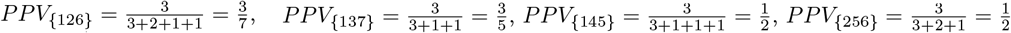, Thus 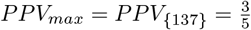.
2. The complex contains more than three subunits. Without loss of generality, consider a protein complex consisting of four subunits. Similarly to the above, it is clear that there is a chance that PPV will be maximized when this complex contains a complex from the reference set. Let the complex in the prediction set be *PC*_1_ and the corresponding complex in the reference set contained by *PC*_1_ be *RC*_1_. Since only complexes containing three subunits exist in the reference set, for *PC*_1_, the increase of subunit compared with *RC*_1_ does not increase the maximum matching number of subunits in the numerator of *PPV*. On the other hand, additional subunits of *PC*_1_ may exist in other complexes from the reference set, which will increase the denominator of *PPV*. This means 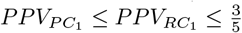. When there are more than one complex in the prediction set, we refer to it as *PC* _1_, *PC* _2_, … *PC* _*m*_, where the complex with the largest *PPV* is denoted as *PC* _*_. Let 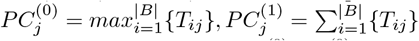. In contrast to Eq. 1, we have 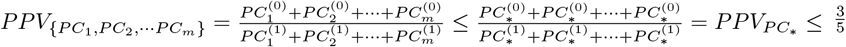□.

